# PRC2 clock: a universal epigenetic biomarker of aging and rejuvenation

**DOI:** 10.1101/2022.06.03.494609

**Authors:** Mahdi Moqri, Andrea Cipriano, Daniel Nachun, Tara Murty, Guilherme de Sena Brandine, Sajede Rasouli, Andrei Tarkhov, Karolina A. Aberg, Edwin van den Oord, Wanding Zhou, Andrew Smith, Crystal Mackall, Vadim Gladyshev, Steve Horvath, Michael P. Snyder, Vittorio Sebastiano

## Abstract

DNA methylation (DNAm) is one of the most reliable biomarkers for aging across many mammalian tissues. While the age-dependent global loss of DNAm has been well characterized, age-dependent DNAm gain is less specified. Multiple studies have demonstrated that polycomb repressive complex 2 (PRC2) targets are enriched among the CpG sites which gain methylation with age. However, a systematic whole-genome examination of all PRC2 targets in the context of aging methylome as well as whether these associations are pan-tissue or tissue-specific is lacking. Here, by analyzing DNAm data from different assays and from multiple young and old human and mouse tissues, we found that low-methylated regions (LMRs) which are highly bound by PRC2 in embryonic stem cells gain methylation with age in all examined somatic mitotic cells. We also estimated that this epigenetic change represents around 90% of the age-dependent DNAm gain genome-wide. Therefore, we propose the “PRC2 clock,” defined as the average DNAm in PRC2 LMRs, as a universal biomarker of cellular aging in somatic cells. In addition, we demonstrate the application of this biomarker in the evaluation of different anti-aging interventions, including dietary restriction and partial epigenetic reprogramming.

## Introduction

DNA methylation (DNAm) is the epigenetic modification most strongly associated with aging ^1^. Numerous studies have identified examples of specific CpGs whose methylation levels are strongly correlated with age, which have been used to build epigenetic clocks ^2^. In parallel, several studies have demonstrated that there is a global increase in epigenetic entropy as a result of the loss of epigenetic information, which occurs when regions that are predominantly hypo- or hyper-methylated at birth gradually transition to a state of partial methylation ^3,4^. CpG-based clock methods suffer from limited mechanistic interpretability as the CpGs are chosen based purely on their correlation with age ^2,5^. On the other hand, measures of epigenetic entropy assume that the methylation states trend towards partial methylation and increased randomness. This basic assumption does not hold for sites most highly correlated with age used in epigenetic clocks as they transition from one methylation state toward the other ^6^. We have recently demonstrated that during aging, mitotic somatic cells undergo a global loss of DNAm particularly in heterochromatic gene-poor regions ^7^. To extend our previous work, in this study, we perform an extensive analysis of the age-dependent gain of DNAm. A relatively small fraction of mammalian genomes, best exemplified by broad CpG islands (CGI) and certain non-CGI sites, lacks DNA methylation ^8^. They are known as low-methylated regions ^9^, non-methylated islands ^10^, DNAm valleys ^11^, DNAm canyons ^12^, and DNAm nadir ^13^, in the existing literature and are highly enriched in regulatory regions ^12^. LMRs at actively transcribed promoters are proposed to remain devoid of DNAm through the formation of DNA-RNA hybrid structures that protect the DNA from methyltransferase activity ^14^. LMRs at repressed promoters are believed to actively maintain hypomethylation through the methylcytosine dioxygenase TET1 while facilitating the binding of PRC2 ^15^. The biochemical antagonism between DNA methylation and PRC2 action has been well-documented and found to be ^16–19^. PRC2 binding regions are enriched for cancer-associated hypermethylation ^20–22^. These regions are also enriched in age-associated hypermethylation in cancer ^23^ and in normal adults ^24^. While previous studies have identified the enrichment of PRC2 targets within the age-dependent methylome ^23,25–30^, they are mainly based on a selected number of CpG sites or a small number of samples. In addition, the commonly used microarray assays in previous studies only cover a small fraction of all CpGs genome-wide (1-3 percent of all CpGs in Illumina BeadChip arrays, Illumina.com 2022). This is insufficient for assessing the association between PRC2 binding and age-dependent gain of methylation genome-wide. Here, we examine genome-wide age-dependent changes in DNAm by using multiple bulk and single-cell Whole-Genome Bisulfite Sequencing (WGBS), Reduced-Representation Bisulfite Sequencing (RRBS), and HumanMethylation450 BeadChip (HM450) microarrays datasets of young and old samples from different human and mouse tissues. We then propose the PRC2 clock as a biologically informed approach to measure aging, based on the progressive gain of DNAm at low-methylated regions which are highly bound by PRC2 (PRC2 LMRs) genome-wide. This method is assay-agnostic, robust to site-specific variability and noise, and identifies age-dependent changes at the level of genomic regions, such as chromosomes. In contrast to our recent work on global cell-agnostic loss of DNAm with aging in gene-poor heterochromatin regions, this work identifies genome-wide gain of methylation at LMRs with aging.

## Results

### 1. Low-methylated regions are common across different cell types

Many of the region’s most commonly observed in the studies of the aging methylome, such as ELOVL2 promoter, are located at low-methylated regions (LMRs) in the tissue of interest. In addition, these same LMRs are also lowly methylated in other cell types and tissues, including human embryonic stem cells (hESCs). To systematically examine this pattern, we analyzed WGBS data from young (neonatal) and old (centenarian) human CD4+ T cells, young and old human epidermis (3 young and 3 old pooled epidermis samples from sun-exposed skin), and hESCs (H1 cell line) by identifying and comparing LMRs within each sample. From this analysis, we observed a significant overlap in LMRs in the 3 cell types analyzed. We found that 90% of LMRs in T-cells and 67% of LMRs in the epidermis are overlapping (methods, Fig. 1a). In addition, 91% of the T-cell/epidermis common LMRs are also lowly methylated in hESCs (Fig1. a). In agreement with previous studies, we also observed that these regions are highly bound by PRC2 core subunits, EZH2 and SUZ12 in hESCs (see ELOVL2 promoter, for example, Fig1. b). This association seems to explain a major component of the aging methylome as the regions bound by PRC2 represented 91-92 % of all LMR regions greater than 100 bps which gain DNAm (delta DNAm > 0.05) in T cells and epidermis (Fig1 c). A similar trend was also observed in mouse by analyzing WGBS data from young versus old hepatocytes, but to a lower extent, e.g., for a lower delta DNAm, as the age-dependent DNAm gain seems to be smaller in mouse (see next section for more details).

**Fig 1.**
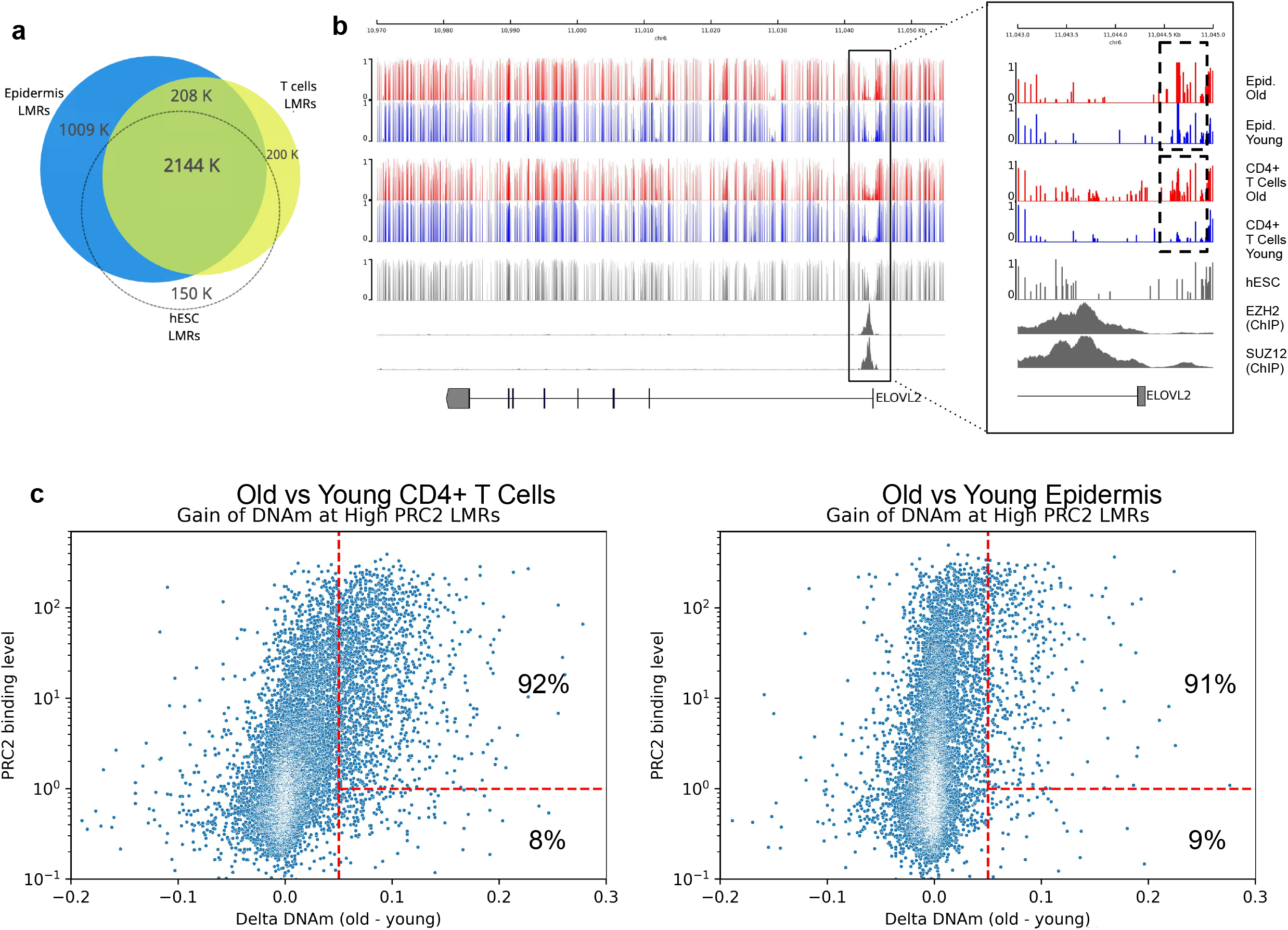
Low-methylated regions are common across different cell types. **a**, Venn diagram showing the overlap between the low methylated regions (LMRs) identified in: epidermidis (GSE52972), CD4+ T cells (GSE31263), and hESCs (H1, GSE16256). **b**, Genome browser visualization of ELOVL2 gene locus showing the DNA methylation tracks of: Old and young Epidermidis (GSE52972), old and young CD4+ T cells (GSE31263) and the ChIP-seq coverage of SUZ12 and EZH2 in hESCs from ENCODE. **c**, Dotplot showing the correlation between PRC2 binding levels and the Delta DNAm between young and old CD4+ T cells (left panel) and epidermidis (right panel). LMRs with PRC2 binding level greater than 1 (see methods) represent 91-92% of regions which gain DNAm.

### 2. PRC2 targets gain DNA methylation by age

Prompted by our previous findings (Fig. 1) we examined the association between PRC2 binding in hESCs and DNA methylation genome-wide and across species by comparing DNAm levels within all LMRs between young and old samples. WGBS data from human CD4+ T cells (aged 18 to 100 years; cord blood has age 0), human epidermis (12 samples from 3 young and 3 old subjects, each with sun-exposed and sun-protected skin), as well as mouse hepatocyte samples (4 samples aged 2 months and 4 samples aged 22 months), were analyzed (Fig. 2a). While the overall DNAm level during aging is generally decreased (Fig. 1a left panels), as we previously showed ^7^, we observed that the low-methylated regions which are highly bound by PRC2 (PRC2 LMRs, methods) gain methylation with age in all the samples analyzed. (Fig 2a, right panels). This robust and consistent age-dependent gain of methylation pattern allowed us to appreciate the trend also at the chromosomal level (Fig 2b). Interestingly, by comparing epidermis samples from 6 young (18 to 25 years) and 6 old (74 to 83 years) individuals, we first confirmed the same trend in DNAm gain at PRC2 LMRs observed in human T-cells (Fig. 2a). We also observed that while no differences were found in young sun-exposed vs sun-protected samples, old samples from sun-exposed epidermis were significantly more methylated in these regions, compared to sun-protected epidermis samples from the same individuals (Fig 2c). This evidence suggests that this phenomenon is not only strongly dependent on the biological age but also on the external environment which may play a pivotal role in accelerating such biological effects.

**Fig 2.**
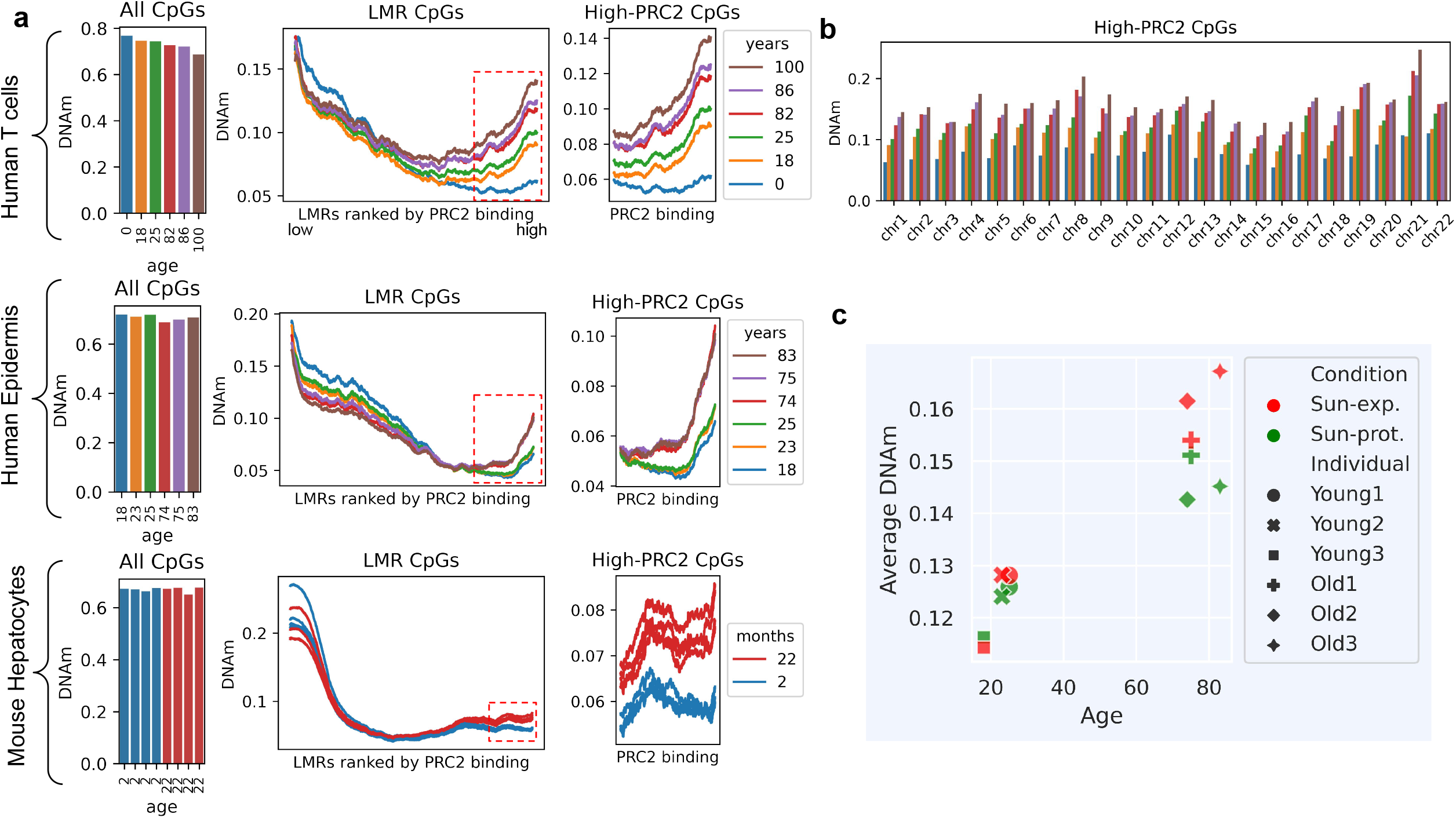
PRC2 targets gain DNA methylation by age. **a**, Average DNAm levels at CpGs across the whole genome (left panel) and at the low methylated regions (LMRs) ranked by the level of their PRC2 binding in hESCs (right panels). Data were obtained by analysing WGBS datasets from human CD4+ T cells (6 samples from 18 to 86 years old, GSE79798 and 2 samples at 0 and 100 years, GSE31263), human epidermis (12 samples from 6 different ages, each age including a sun-exposed and sun-protected sample, that were averaged for the analysis, GSE52972), and mouse hepatocytes (8 samples, 4 2-months vs 4 22-months (GSE89274). PRC2 binding was evaluated by using EZH2 ChIP-seq data from ENCODE in H1 and H9 cell lines. One outlier sample is removed (methods) **b**, Average DNAm levels at LMRs with high PRC2 binding showed for each autosomal chromosome. **c**, Average DNAm levels at LMRs with high PRC2 binding. Data were produced by analyzing the same WGBS datasets from human epidermis shown in **a.**

### 3. PRC2 clock is measurable using different assays

Encouraged by the compelling results obtained from WGBS data, we asked whether this effect was also measurable using different targeted assays. To answer this question, we extended our analysis to 4,518 HM450 microarray blood samples of different ages (13 to 100 years old), from four large publicly available DNA methylation datasets (Table 1). While an overall loss in DNAm was not observed given the low coverage of the arrays (1.5 % of all CpGs), by comparing all low-methylated CpGs and all CpGs within PRC2 LMRs, we observed a consistent gain of methylation within PRC2 LMRs across all four datasets (Fig 2a). We also analyzed a large dataset of RRBS samples from 6 different tissues (adipose, kidney, muscle, lung, liver, and blood, Table 1) from young and old mice. By comparing all low-methylated CpGs and all CpGs within PRC2 LMRs, we observed a consistent gain of methylation within PRC2 LMRs across all the tissues (Fig 2b), suggesting that the gain of methylation at PRC2 sites is evolutionarily conserved in mammals and can be captured using different DNA methylation assays. Notably, similar results were also obtained by analyzing DNAm levels at PRC2 LMRs in mouse liver by using single-cell WGBA (scWGBA) data (Fig 2c), demonstrating the robustness of our method with noisy, low coverage data

### 4. PRC2 clock as a biomarker of rejuvenation treatments

The robust correlation we observed between DNAm at PRC2 LMRs and physiological aging prompted us to examine if it could be used as a biomarker for rejuvenation treatments. To test this hypothesis, we examined the differences in methylation levels on PRC2 LMRs in response to several rejuvenation interventions. We analyzed two publicly available genome-wide methylation sequencing datasets of aging interventions in mouse liver (WGBS) and blood (RRBS) and calculated methylation levels at PRC2 LMRs for all samples (Table 1). In mouse liver, we observed old mice subjected to long-term caloric restriction (CR) had slightly lower methylation levels compared to old control mice (significant only in one-tailed t-test after removing outliers, p-value= 0.039, details in the methods) while no significant decrease in methylation was observed in mice treated with rapamycin (Fig 4a). This suggests that the anti-aging effects of CR could be associated with amelioration of methylation gain at PRC2 LMRs, while it is less likely that rapamycin induces epigenetic rejuvenation via the demethylation of highly methylated PRC2 LMRs. Similarly, we observed lower methylation in CR mice compared to control mice, across 195 RRBS blood samples (Fig. 4b). We also tested if the PRC2 clock could also be used to examine partial epigenetic reprogramming, a relatively new rejuvenation treatment that has become increasingly popular recently ^31–33^. To study the rejuvenation effects of epigenetic reprogramming, we examined DNAm changes in a rejuvenation treatment of mouse skin through partial reprogramming via Yamanaka factors (OSKM^33^, profiled *in vivo* using Infinium Methylation EPIC DNA methylation data (<u>GSE190665</u>). Interestingly we found that both fully and partially reprogrammed samples had significantly lower PRC2 LMRs methylation levels compared to the control samples (p-value 0.03, Welch’s t-test for unequal variance, Fig. 4c), suggesting that the age-dependent gain of methylation is reversible and can be used to assess the levels of epigenetic rejuvenation. Taken together, these data clearly demonstrate that our PRC2 clock is a reliable, agnostic, and universal method that, in addition to measuring biological aging, can also quantify the effect of rejuvenation on epigenetic aging.

### 5. PRC2 LMRs gain DNA methylation by mitosis

Given the known relationship between cell replication and the loss of DNA methylation ^7^, we decided to examine the effect of prolonged cell replication in fully differentiated cells on DNAm gain within PRC2 LMRs. We assessed this effect in WGBS data from cancer and healthy samples for 8 different cancer types from The Cancer Genome Atlas (TCGA, see Zhou et al. 2018 for details, Fig3 a). First, we confirmed the general loss of DNAm at CpGs sites genome-wide (Fig 5a, left panels) as expected. In addition, we also found that similar to the aging methylome, high PRC2 regions gain DNAm in all cancer types. On the other hand, oligodendrocytes, which are post-mitotic in undamaged brains ^34^, did not show significant DNAm gain in old individuals compared to the young (Fig. 3b). While our previous HM450 microarray studies of brain samples demonstrated that PRC2 targets are more likely to gain DNAm compared to non-targets ^26^, we believe that this effect is amplified in mitotic cells. To confirm the gain of DNAm is linked to cell replication we analyzed a WGBS dataset of a primary human neonatal foreskin fibroblast at 5 different passages (from passage 4 to passage 34, see ^35^ for details). We found that the average DNAm within PRC2 LMRs across these time points showed a very high correlation with passage number (Fig. 3c) providing additional evidence that cell replication of somatic cells highly contributes to the observed gain of methylation.

**Fig 3.**
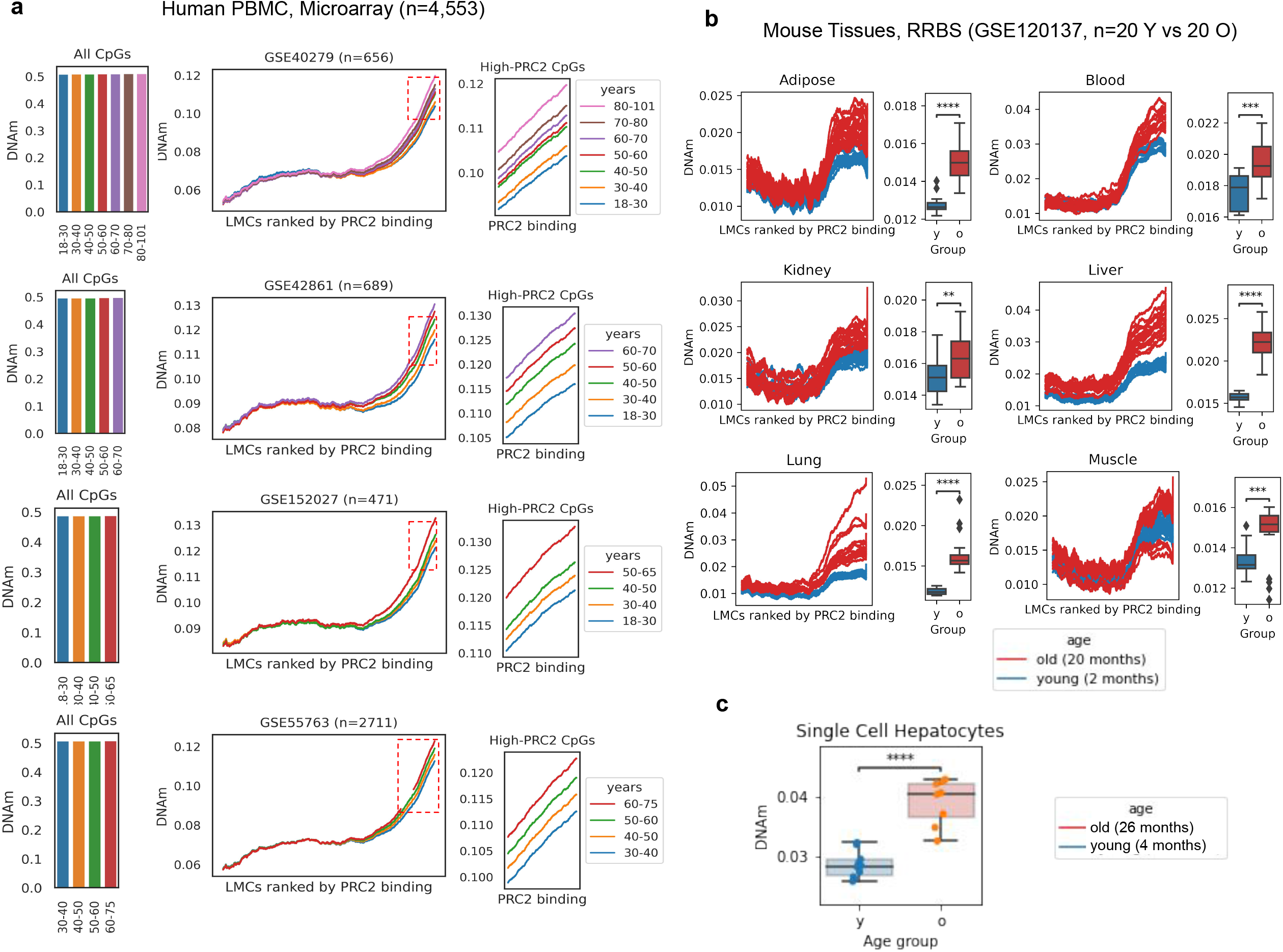
Gain of DNAm at PRC2 targets is measurable using different assays. **a.** Average DNAm levels at CpGs across the whole genome (left panel) and at the LMRs ranked by the level of their PRC2 binding in hESCs (right panels). Data were obtained by analyzing DNA methylation microarray data from human PBMC samples. Data from four different HM450 datasets (GSE40279, GSE42861, GSE152027, and GSE55763) are shown in each panel. PRC2 binding was estimated by using EZH2 and SUZ12 ChIP-seq data from ENCODE in H1 and H9 cell lines (methods). **b.** Average DNAm levels in different mouse tissues at low methylated CpGs (LMCs) ranked by the level of their PRC2 binding in hESCs (left panel). The right panel shows the same data represented as averaged. Data were obtained from a RRBS dataset (GSE120137). One outlier mouse was removed from all analyses (see methods). **c.** Average DNAm levels at low methylated LMCs with high PRC2 binding in hESCs, in mouse liver from scWGBS dataset (SRP069120). One outlier was removed from each category according to Trap et al. 2021, methods). p-value levels: * < 0.05, **: < 0.01, ***: < 0.001, ****: < 0.0001.

**Fig 4.**
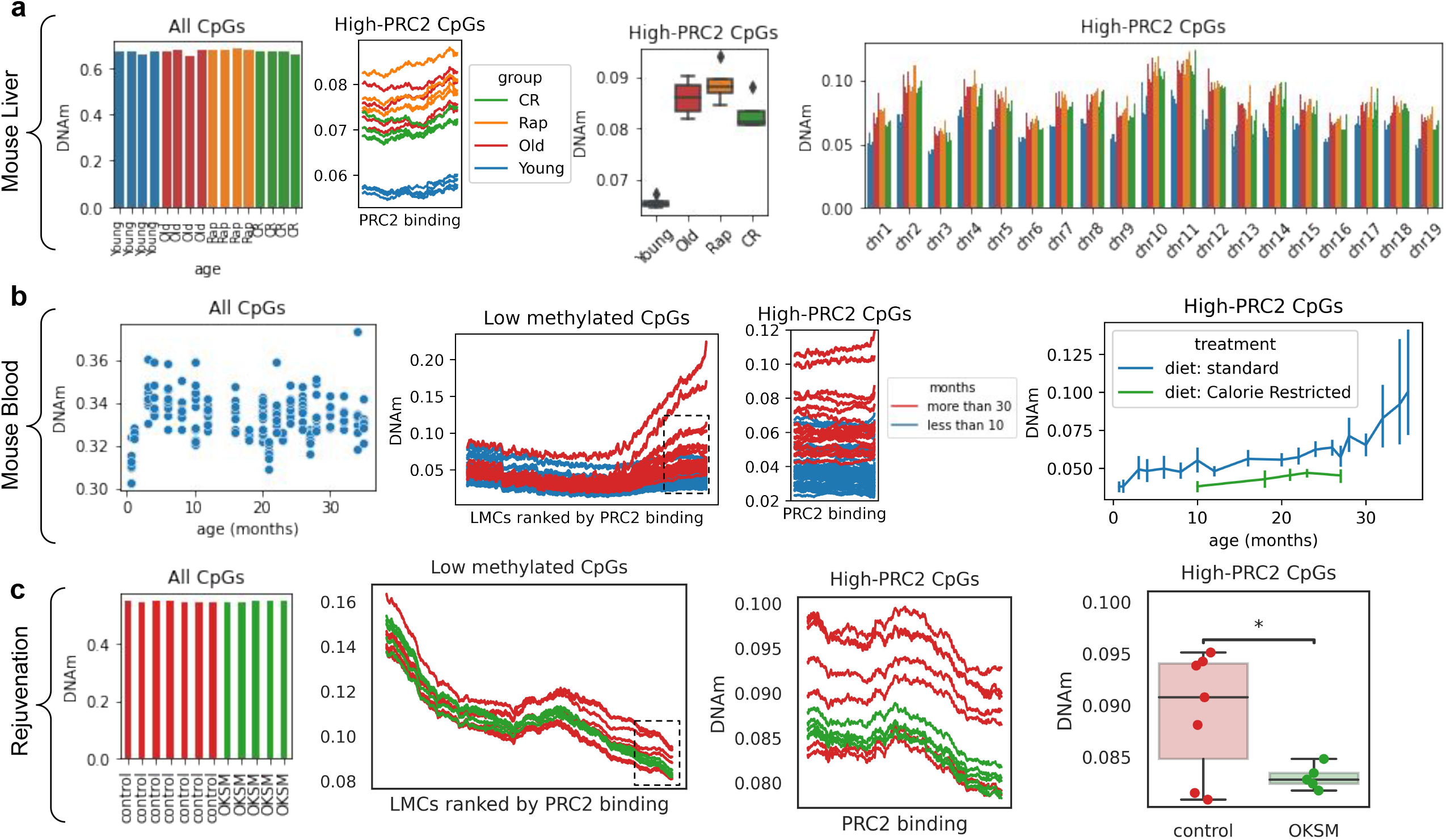
PRC2 targets DNAm as a marker of rejuvenation. **a.** Average DNAm levels at CpGs across the whole genome (left panel), at the LMRs showing a high PRC2 binding in hESCs (middle panel), in liver samples from young, old, caloric restricted (CR), and rapamycin treated (Rap) mice. The box-plot on the right, shows the averaged DNAm levels between the 4 biological replicates. (WGBS dataset GSE89274, n=16) **b.** Average DNAm levels at CpGs across the whole genome (left panel) and at the LMRs ranked by the level of their PRC2 binding in hESCs (middle panels), average DNAm levels in High PRC2 LMCs at different ages in blood samples from ctrl and caloric restricted mice (RRBS dataset GSE80672, n=110). **c.** Average DNAm levels at CpGs across the whole genome (left panel), and at the LMRs ranked by the level of their PRC2 binding in hESCs (middle panels (middle and left panel) from DNA methylation microarray long-term partial reprogramming experiments (dataset GSE190665, n=12).

**Fig 5.**
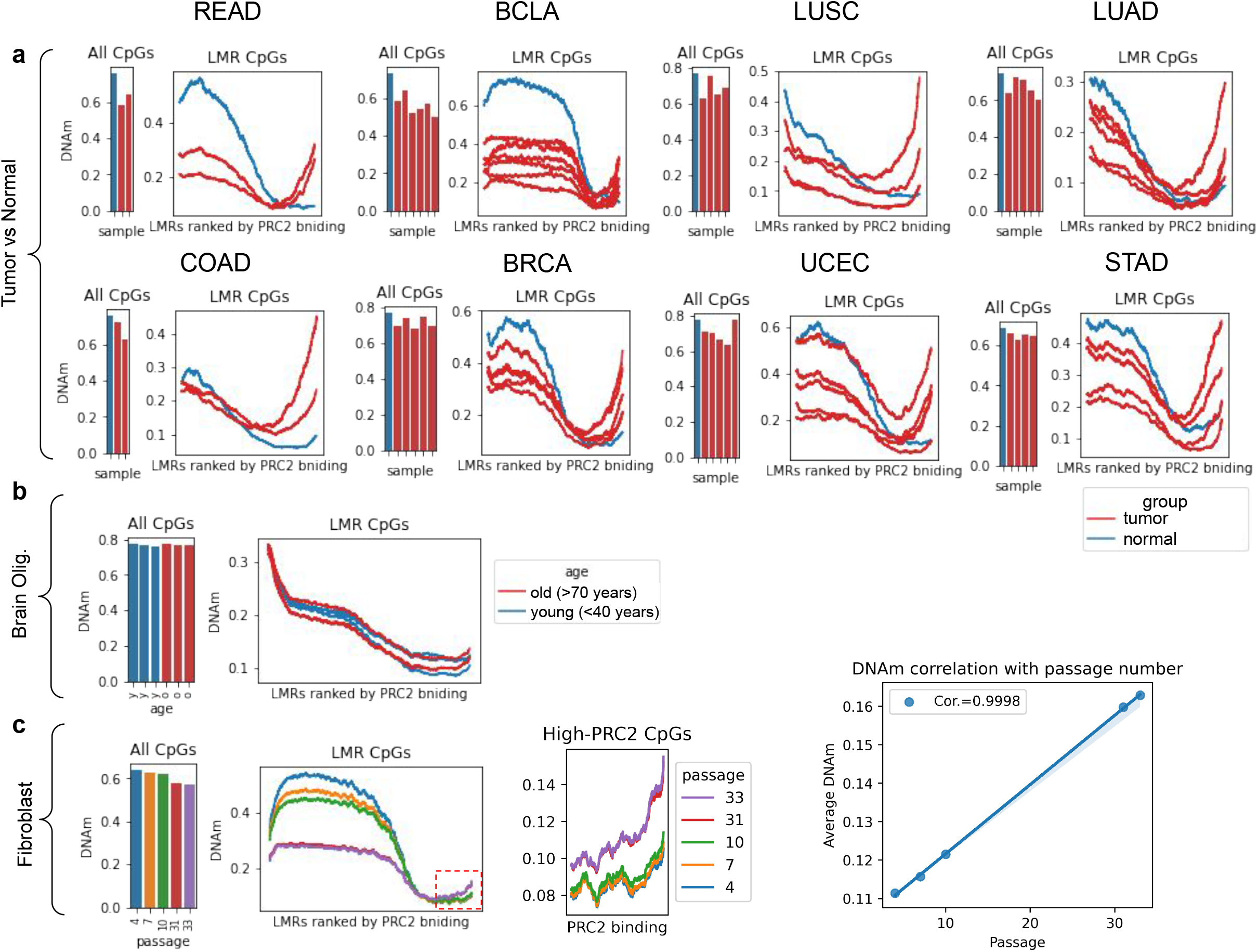
PRC2 targets gain DNA methylation by mitosis. **a**, Average DNAm levels at CpGs across the whole genome (left panel) and at LMRs ranked by the level of their PRC2 binding in hESCs (right panel) from different healthy vs breast cancer samples. Data were obtained from TCGA WGBS datasets **b**, Average DNAm levels at CpGs across the whole genome (left panel) and at LMRs ranked by the level of their PRC2 binding in hESCs (right panel) from young vs old human oligodendrocytes. Data were obtained from WGBS GSE107729 datasets. **c**, Average DNAm levels at CpGs across the whole genome (left panel) and at LMRs ranked by the level of their PRC2 binding in hESCs (middle panel), correlation between average DNAm at high-PRC2 LMRs and the number of passages in *in vitro* cultured fibroblasts (right panel). Data were obtained from GSE79798 WGBS datasets.

## Discussion

This study takes a significant step toward addressing two known major challenges in epigenetic studies of aging: a comprehensive genome-wide study of the aging methylome and the identification of interpretable epigenetic markers ^5,36^. In contrast to previously developed epigenetic aging biomarkers, our approach is biologically informed and does not rely on “black-box” predictive models that are challenging to interpret. Through the whole-genome examination of PRC2 LMRs, our model highlights a specific cellular process associated with aging. One possible explanation for why PRC2 LMRs have low methylation in young cells is to reduce the risk of somatic genetic mutations by protecting them from spontaneous deamination of methylated cytosines ^37^. Spontaneous deamination is one of the primary mechanisms through which oncogenic mutations arise ^38^, so there may be a selective pressure to protect critical CpGs from deamination in younger individuals that is lost with aging. In support of this hypothesis, recent cancer epigenome studies have demonstrated significant DNAm gain at PRC2 LMRs ^39,40^. Further experiments are needed to examine the direction and causality of this effect. Previous studies in humans and other mammalian species have shown that target sites of PRC2 gain methylation with age ^23,25–30^, although this was identified as an enrichment among a small number of CpGs rather than a global survey of PRC2 targets. Two different biological processes have previously been suggested to contribute to the age-dependent gain of DNAm within PRC2 LMRs. First, cells respond to double-strand breaks by translocating chromatin modifiers (e.g. DNMT1) to the site of the break, often from CpG-poor to CpG-rich regions which are enriched for PRC2-LMRs. This in turn could result in an increase in DNAm within these regions ^36,41^. Second, PRC2 binding to DNA is suggested to protect unmethylated CpG-rich regions from becoming methylated and age-related degradation of the PRC2 machinery could result in the decline of PRC2 function in identifying and/or binding its targets, consequently resulting in de novo methylation of these regions via DNMT3A and DNMT3B ^36,42^. Here, we suggest a third potential contributor to the effect. As most PRC2 LMRs contain gene promoters which are silent or lowly expressed in somatic cells, these regions might afford to gain marginal DNAm without being immediately deleterious to cells. However, these gradual age-dependent epigenetic changes might contribute to the functional dysregulation associated with aging. Ongoing mechanistic studies of the interactions between methyltransferases and PRC proteins will give further insight into the maintenance of DNAm at PRC2 LMRs and how it might be dysregulated during aging ^43^. Future studies in non-renewing cell types such as neurons will be needed to determine if the mechanism that associates PRC2 LMRs with aging differs from dividing cells. Similarly, measuring PRC2 LMR methylation in adult stem cells at different states of exhaustion from the same individual could provide insight into the heterogeneity of epigenetics within a single tissue. In summary, our results demonstrate that we have developed a new, robust method to assess the aging methylome using biologically informed features. Our method does not require training, does not rely on specific CpGs and works across species and methylation assays. Most importantly, it can assess the aging of primary cells both *in vivo* and *in vitro* and measure the effect of epigenetic rejuvenation on cells and tissues.

## Acknowledgments and Fundings

V.S. is supported by the MCHRI Woods Family Endowed Scholarship in Pediatric Translational Medicine (Stanford Maternal & Child Health Research Institute), by the Breakthrough in Gerontology Award (BIG Award, AFAR/Glenn Foundation) and by the Milky Way Research Foundation. A.C. is supported by the DiGenova Postdoc Seed Grant (Stanford University). M.M. is funded by NIH T15 in Biomedical Informatics and NIH T32 Aging Research at Stanford.

## Conflict of Interest

V.S. is a co-founder, SAB Chairman, and a shareholder of Turn Biotechnologies. S.H. is a founder of the non-profit Epigenetic Clock Development Foundation

## Materials and Methods

### External WGBS and RRBS data

Whole Genome Bisulfite Sequencing reads were downloaded from the Sequence Read Archive (SRA). Reads were preprocessed by removing adapters using *<u>(Babraham Bioinformatics - Trim Galore)</u>* version 0.6.7 and subsequently mapped to their corresponding reference genome (either hg38 or mm10) using *abismal* version 2.0.0 ^44^. The output SAM files were analyzed using methpipe version 5.0.1 ^45^. Paired-end reads were merged using the *format_reads* tool, after which reads were sorted by chromosome coordinates using *samtools sort* ^46^. PCR duplicates, defined as identical reads that map to the same location, were removed using the *duplicate-remover* tool in methpipe. Individual CpG methylation levels were quantified from SAM files using *methcounts* and summary statistics on CpG coverage, depth, and average methylation were quantified using the *levels* program.Methylation levels of symmetric CpGs in opposite strands of the reference were merged using the *symmetric-cpgs* program. Low-methylated regions were identified from symmetric CpG methylation levels using the *hmr* program with default parameters. UCSC genome browser tracks for symmetric CpGs were created using the *<u>wigToBigWig</u>* program. LMRs were ranked based on the average PRC2 binding levels (see ChIP-Seq section). The horizontal axis in all LMR figures shows ranked PRC2 regions from lowest binding (left) to highest binding (right). In order to reduce the noise and visualize the trends in DNAm changes, sliding windows and smoothing are used and the average DNAm is calculated and displayed within each sliding window.

### HM450 analysis

All HM450 datasets were downloaded from GEO (GSE152027, GSE40279, GSE42861, and GSE55763). No additional normalization of methylation levels was performed.

### Outliers

Among the 8 human WGBS CD4+ T samples, one adult sample was removed since its overall methylation was outside the range between the neonatal and centenarian samples. Among the 40 mouse RRBS samples, one mouse had significantly higher methylation levels across different tissues and was removed from the analyses. Among scWGBS hepatocyte samples, following the analyses by Trapp. et al. 2021, one sample was removed from each of the young and old groups. A list of all datasets with outliers is available in Table 1.

### ChIP-Seq

EZH2 and SUZ12 binding regions and levels were extracted from ENCODE ChIP-seq data from H1 human ESC cell line publicly available under accession numbers SR000ASY and SR000ATS and from mouse embryonic cells lines (E14) publicly available under accession numbers GSM2472741 and GSM3243624. PRC2 bound regions are identified using p-values from the Encode pipeline (p-value to reject the null hypothesis that the signal at that location is present in the control^47^). High-PRC2 LMRs are selected based on the top 1000 LMRs with the lowest average p-values from EZH2 and SUZ12 Chip-Seq data.

### Code availability

All of our scripts to analyze the data, produce the figures, and generate the final PRC2 LMRs are freely accessible on Github at https://github.com/moqri/aging (will be made public for publication)

## Data availability

Our fibroblast WGBS raw data are available in the Sequence Read Archive (SRA) under accession number PRJNA804539 (public upon publication)

## Notes

https://www-ncbi-nlm-nih-gov.ezproxy.u-pec.fr/geo/query/acc.cgi?acc=GSE52972

https://www.ncbi.nlm.nih.gov/geo/query/acc.cgi?acc=GSE31263

https://www.ncbi.nlm.nih.gov/geo/query/acc.cgi?acc=GSE16256

https://www.ncbi.nlm.nih.gov/geo/query/acc.cgi?acc=GSE79798

https://www.ncbi.nlm.nih.gov/geo/query/acc.cgi?acc=GSE89274

https://www.ncbi.nlm.nih.gov/geo/query/acc.cgi?acc=GSE152027

https://www.ncbi.nlm.nih.gov/geo/query/acc.cgi?acc=GSE40279

https://www.ncbi.nlm.nih.gov/geo/query/acc.cgi?acc=%20GSE42861

https://www.ncbi.nlm.nih.gov/geo/query/acc.cgi?acc=GSE55763

https://www.ncbi.nlm.nih.gov/geo/query/acc.cgi?acc=GSE120137

### Competing Interest Statement

V.S. is a co-founder, SAB Chairman and shareholders of Turn Biotechnologies.
SH is a founder of the non-profit Epigenetic Clock Development Foundation

